# Identification of COVID-19 and COPD common key genes and pathways using a protein-protein interaction approach

**DOI:** 10.1101/2021.10.28.466298

**Authors:** Thiviya S. Thambiraja, Kalimuthu Karuppanan, Gunasekaran Subramaniam, Suresh Kumar

## Abstract

Coronavirus disease (COVID-19) is an extremely contagious and cognitive disease that could cause immense hypoxemia. The rise in critically ill patients in epidemic regions has put enormous pressure on hospitals. There is a need to define extreme COVID-19 clinical determinants to optimize clinical diagnosis and the management system is strong. Chronic obstructive pulmonary disease (COPD) is linked to a rapidly increasing risk of death rates in population pneumonia. In this research, a network of protein-protein interaction (PPI) was developed using constructed datasets of COVID-19 and COPD genes to define the interrelationship between COVID-19 and COPD, how it affects each other, and the genes that are responsible for the process. The PPI network shows the top 10 common overlapping genes, which include IL10, TLR4, TNF, IL6, CXCL8, IL4, ICAM1, IFNG, TLR2, and IL18. These are the genes that COVID-19 and high-risk COPD patients are known to be expressed. These important genes shared by COVID-19 and COPD are involved in pathways such as malaria, African trypanosomiasis, inflammatory bowel disease, Chagas disease, influenza, and tuberculosis.

## INTRODUCTION

At the end of 2019, a novel coronavirus named SARS-CoV-2 emerged in the Chinese city of Wuhan, sparking an epidemic of rare viral pneumonia. Since it was designated a pandemic, this unusual coronavirus infection, named coronavirus disease 2019 (COVID-19), has spread fast around the world [1]. As of October 28th, 2021, there were 245,842,215 reported cases and 4,988,705 confirmed death cases. [2] According to the Centers for Disease Control and Prevention (CDC), the risk of having severe COVID-19 illness, which may result in hospitalisation and death, increases with age. Indeed, COVID-19 has been implicated in eight out of ten deaths among persons 65 years of age and older. Additionally, those 65 years of age and older are more likely to have COPD; although infection with COVID-19 is rare in COPD patients, those with COPD who are infected with the virus have a higher mortality risk than those without COPD. Given COVID-19’s life threatening impact on the lung, those with underlying COPD should be concerned. Calculating their increased risk of contracting COVID-19 and associated more severe respiratory symptoms in this pandemic has been problematic for a multitude of reasons. Additionally, differential diagnosis of COVID-19 is challenging since its symptoms may resemble those of chronic and progressive COPD. Consequently, it is vital for COPD patients experiencing an exacerbation to be tested for the virus and to practise social isolation in order to prevent the virus from spreading. COPD is a heterogeneous condition with a broad range of disease severity, frequency of exacerbations, and comorbidities. In November 2020, the Global Initiative for the Management of Chronic Obstructive Lung Disease (GOLD) published its 2021 COPD Management Report. Numerous major updates to the criteria for diagnosing, evaluating, and treating COPD are included in the Report. Despite this, accumulating evidence indicates that COPD may be a risk factor for more severe COVID-19 disease [3].

Additionally, to dig deeper into COPD, it is the third biggest cause of mortality worldwide and is predicted to continue to be a serious public health problem soon. It is characterised by airflow limitations that affect around 10% of persons over the age of 40. Environmental and genetic factors both contribute to the risk. Though cigarette smoke is a major risk factor, only 20% of smokers acquire the condition. Recent genome-wide association analyses identified the COPD-susceptible genomic region. In contrast to cystic fibrosis, which is a monogenic genetic condition, COPD is a polygenic complicated disease [4-6]. COPD patients are more sensitive to co-morbidity due to the activation of the chemokine network in COVID-19 [7].

COVID-19 infection manifests symptoms after an incubation period of around 5.2 days. The time interval between the commencement of COVID-19 adverse effects and mortality varied between six and forty-one days, with a mean of fourteen days. The duration is determined by the patient’s age and the status of his or her immune system. It was shorter among individuals over 70 years of age compared to those under 70. The most common symptoms associated with the start of COVID-19 are fever, cough, and fatigue. Sputum, diarrhoea, haemoptysis, diarrhoea, lymphopenia, and dyspnoea are only a few of the symptoms. Whereas COPD is a chronic inflammatory lung disease that causes the lungs’ airways to become clogged. Symptoms include difficulty breathing, coughing, mucus (sputum) production, and wheezing. It is most often caused by continuous exposure to irritant chemicals or particulate matter, the most common of which being cigarette smoke. The most common causes of COPD are emphysema and chronic bronchitis. These two disorders often overlap and may vary in severity among COPD patients. Bronchitis chronica is an infection of the bronchial tubes, which carry air to and from the air sacs of the lungs (alveoli). Chronic coughing and mucus (sputum) production daily characterise it. Emphysema is a condition in which the alveoli at the end of the lungs’ smallest airways (bronchioles) get damaged because of hazardous exposure to cigarette smoke and other irritating chemicals and particulate matter. While COPD is a progressive chronic illness, it is treatable. With proper medication, most people with COPD may achieve great symptom management and a high quality of life, as well as a lowered risk of acquiring further associated diseases. [1]. COVID-19 and chronic obstructive pulmonary disease have been documented to co-exist in some individuals [8]. Individuals with severe COPD may be at a greater risk of developing major complications from COVID-19, since COVID-19 affects the respiratory system [3]. Present lung injury means fighting off an infection is difficult on the lungs [9]. According to the researchers, who conducted a meta-analysis of seven studies, persons with COPD may have a significantly increased chance of getting severe COVID-19 infections [10]. COPD is associated with a nearly fourfold increased chance of developing significant COVID-19 [11]. Concerning COVID-19, it was discovered that levels of angiotensin-converting enzyme 2 (ACE2), the virus’s known target receptor, increased in COPD patients [12]. Additionally, there is a statistical value indicating that the prevalence of COPD in hospitalised COVID-19 patients was 0.95 percent (95 percent confidence interval [CI]: 0.43 percent -1.61 percent) [13]. Smokers and patients with COPD had higher airway expression of ACE-2, the COVID-19 virus’s entrance receptor. This might account for the higher risk of severe COVID-19 infection in these subpopulations and emphasises the critical role of smoking cessation. [14].

Studying COPD patients who suffering from COVID-19 at the gene function level, will alleviate the disease severity. With the current advancement in bioinformatics, several complex problems are resolved in disease biology and drug discovery [15-18]. In this study, a protein-protein interaction (PPI) network has been constructed by using constructed datasets of COVID-19 and COPD genes to find the interrelationship between COVID-19 and COPD, how it affects each other, and which genes are responsible for the mechanism involved.

## 2 MATERIALS AND METHODS

### 2.1 COVID-19 gene dataset construction

To construct the COVID-19 dataset, the COVID-19 curated geneset has been downloaded from the Comparative Toxicogenomics Database (CTD) (http://ctdbase.org/) [19] DisGeNET (a database of gene-disease associations) (https://www.disgenet.org/) [20] and the study of Gordon et al [21]. All the unique genes were combined to construct the COVID-19 dataset of 5463 genes.

### 2.2 COPD gene dataset construction

To construct the chronic obstructive pulmonary disease (COPD) dataset, the COPD dataset was downloaded from the Comparative Toxicogenomics Database (CTD) and DisGeNET (a database of gene-disease associations) and updated on the COPD gene list from the Yohan Bossé study [6].

### 2.3 Combined gene dataset construction

The overlapping gene sets were combined as one set of overlapping genes. The common overlapping 248 genes COVID-19 and COPD genes were obtained from Venny tools (https://bioinfogp.cnb.csic.es/tools/venny/) [22].

### 2.4 Protein-protein network construction using Cytoscape

To construct a protein-protein interaction (PPI) network, the 248 common overlapping genes were input into the String app [23] in Cytoscape (v 3.8.0) [24] downloaded through the App manager option to build an interaction network. The genes were put through a String protein analysis query and the confidence level of the results was set to 0.40 with 0 extra interactions. Cytohubba [25] downloaded through the App manager option in Cytoscape was then used to calculate all the possible annotations of the interaction network. The Maximal Clique Centrality (MCC) algorithm is used in Cytohubba to calculate the top 10 ranked nodes.

### 2.5 Gene Ontology of top 10 common overlapping genes in COVID-19 and COPD

WebGestalt (WEB-based Gene SeT Analysis Toolkit) http://www.webgestalt.org/ [26] have been used to find the gene enrichments and gene ontology of the top 10 common overlapping genes in COVID-19 and COPD. Homosapiens have been used as the organism of interest, Over-Representation Analysis (ORA) have been selected for the method of interest, and gene ontology selected with biological process non-redundant. The top 10 common overlapping gene symbol have been typed in the text bar and genome protein-coding have been selected for the reference set. Bar charts of biological process categories, cellular components categories, and molecular function categories have been obtained from this analysis.

### 2.6 Gene pathway analysis of top 10 common overlapping genes in COVID-19 and COPD

WebGestalt (WEB-based Gene SeT Analysis Toolkit) http://www.webgestalt.org/ has been used to find the gene pathway of the top 10 common overlapping genes in COVID-19 and COPD. Homosapiens have been used as the organism of interest, Over-Representation Analysis (ORA) have been selected for the method of interest, and for the functional database, the KEGG pathway has been selected. The top 10 common overlapping gene symbol have been typed in the text bar and genome protein-coding have been selected for the reference set. Bar charts of gene pathways have been obtained from this enrichment analysis.

## 3 RESULTS AND DISCUSSION

Based on the data downloaded from the Comparative Toxicogenomics Database (CTD), DisGeNET(a database of gene-disease associations), and the study of Gordon et al, a constructed dataset of 5463 genes COVID-19 were obtained. To create the chronic obstructive pulmonary disease (COPD) dataset, the COPD dataset was obtained from the Comparative Toxicogenomics Database (CTD) and DisGeNET (a database of gene-disease relationships) and updated using the Yohan Bossé study’s COPD gene list containing 296. While the 248 genes shared by COVID-19 and COPD were obtained using Venny tool **(Figure 1)**. The 248 common overlapping genes were put into a query of the String Protein Analysis and the results confidence level was set at 0.40 with 0 additional interactions. Cytohubba available in Cytoscape was then used to measure all potential contact network annotations. The Maximum Clique Centrality (MCC) algorithm is used for determining the top 10 rank nodes in Cytohubba. The top 10 common overlapping genes related to COVID-19 and COPD are IL10, TLR4, TNF, IL6, CXCL8, IL4, ICAM1, IFNG, TLR2, and IL18 **(Table 1). Figure 2** shows the top 10 genes that are common overlapping genes in COVID-19 and COPD highlighted in the PPI network. Figure 3 shows the Gene Ontology of the top 10 common overlapping genes in COVID-19 and COPD. In **Figure 3** which shows the gene ontology analysis, the first bar chart in color red represents the biological process categories, the second bar chart in colour blue represents the cellular components categories and the third bar chart in colour green represents the molecular functions categories. **Figure 4** shows the gene pathway analysis of the top 10 common overlapping genes in COVID-19 and COPD.

**Table 1.**
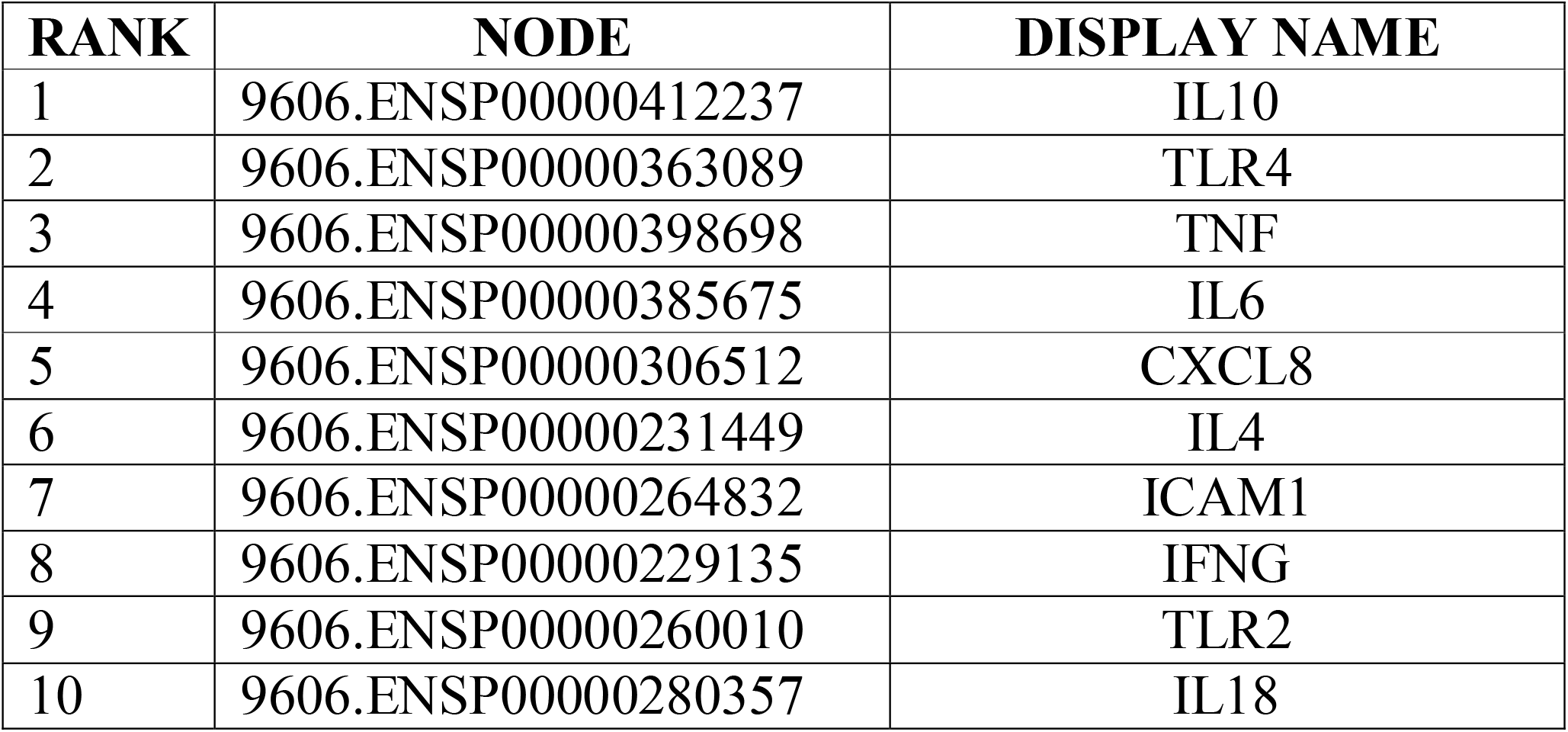
The top 10 genes that are common in COVID-19 and COPD.

| **RANK** | **NODE** | **DISPLAY NAME** |
| --- | --- | --- |
| 1 | 9606.ENSP00000412237 | IL10 |
| 2 | 9606.ENSP00000363089 | TLR4 |
| 3 | 9606.ENSP00000398698 | TNF |
| 4 | 9606.ENSP00000385675 | IL6 |
| 5 | 9606.ENSP00000306512 | CXCL8 |
| 6 | 9606.ENSP00000231449 | IL4 |
| 7 | 9606.ENSP00000264832 | ICAM1 |
| 8 | 9606.ENSP00000229135 | IFNG |
| 9 | 9606.ENSP00000260010 | TLR2 |
| 10 | 9606.ENSP00000280357 | IL18 |

**Figure 1.**
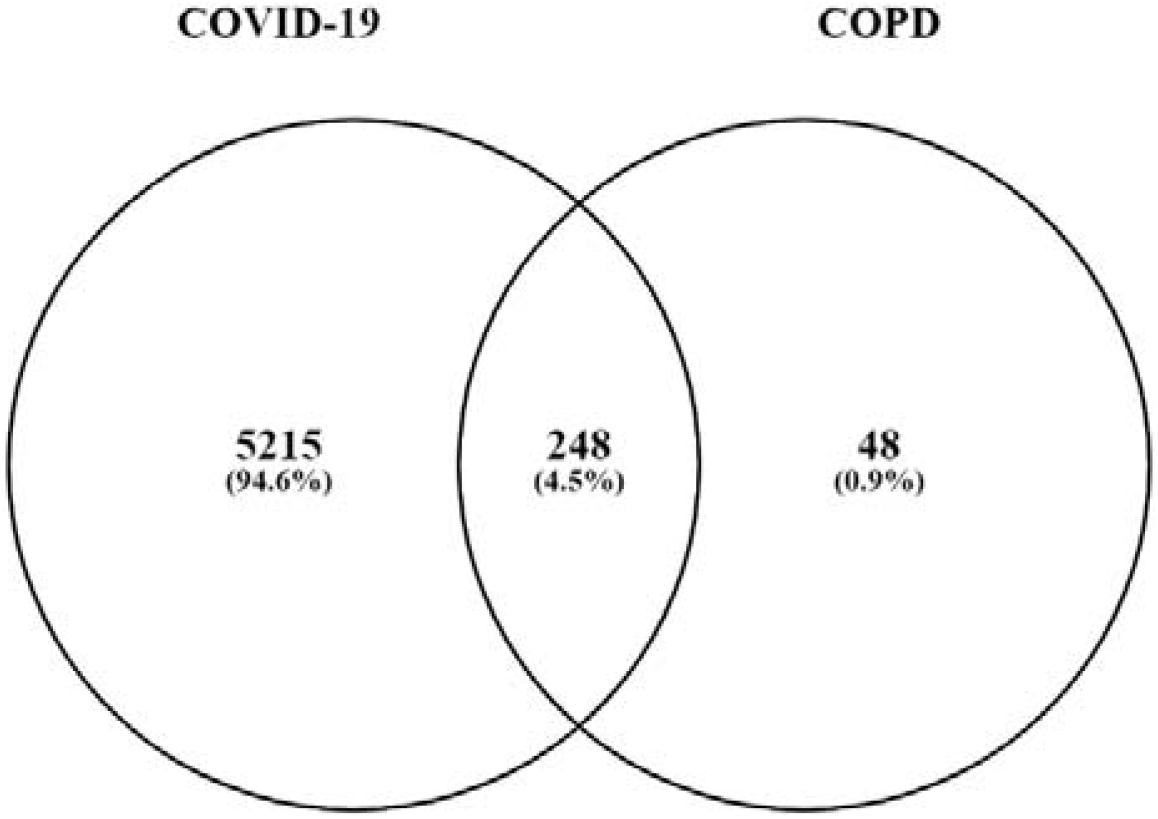
The Venn diagram showing overlapped genes between COVID-19 and COPD.

**Figure 2.**
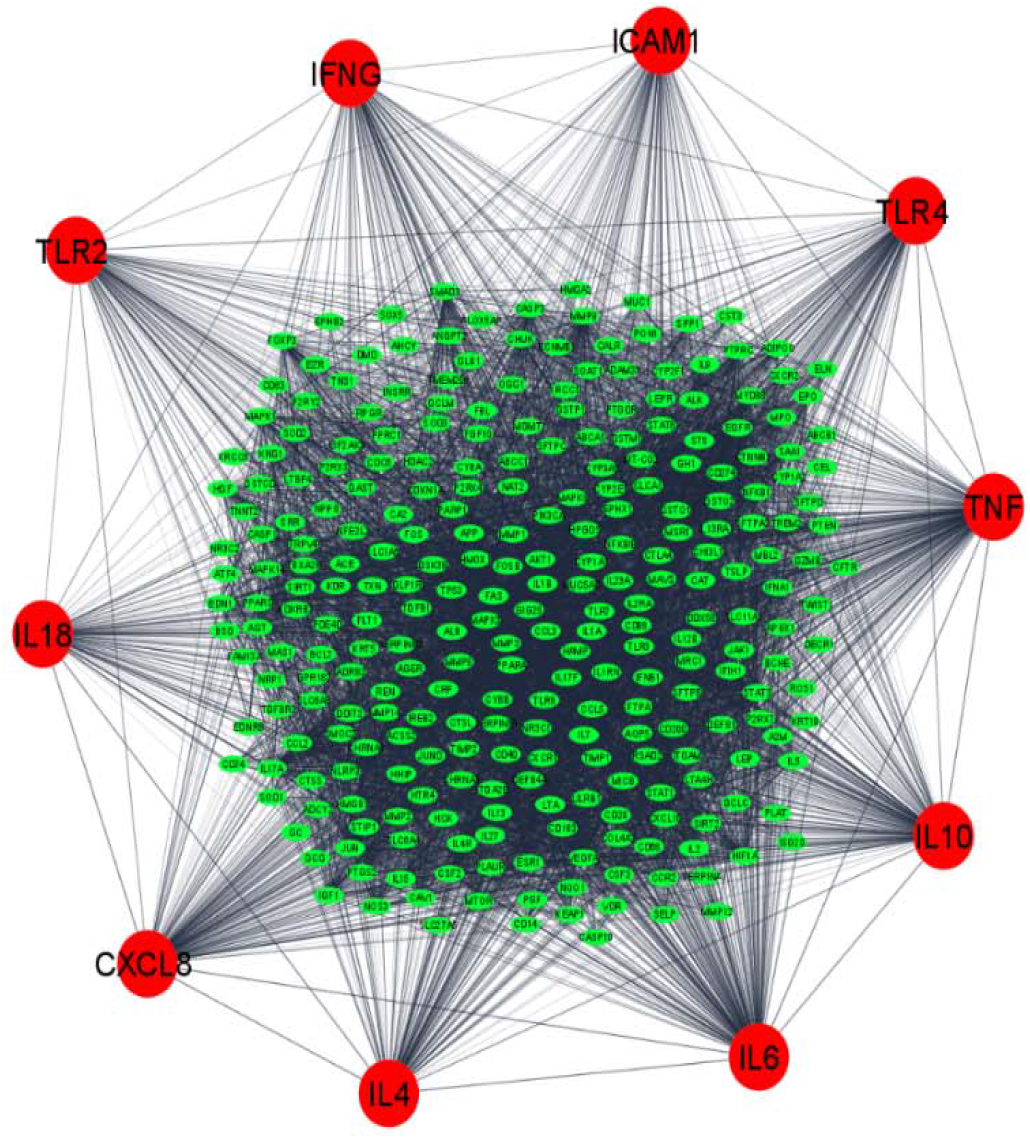
The PPI network of genes that are common overlapping genes in COVID-19 and COPD highlighted in the PPI network. The top 10 hub genes are shown in red colour.

**Figure 3.**
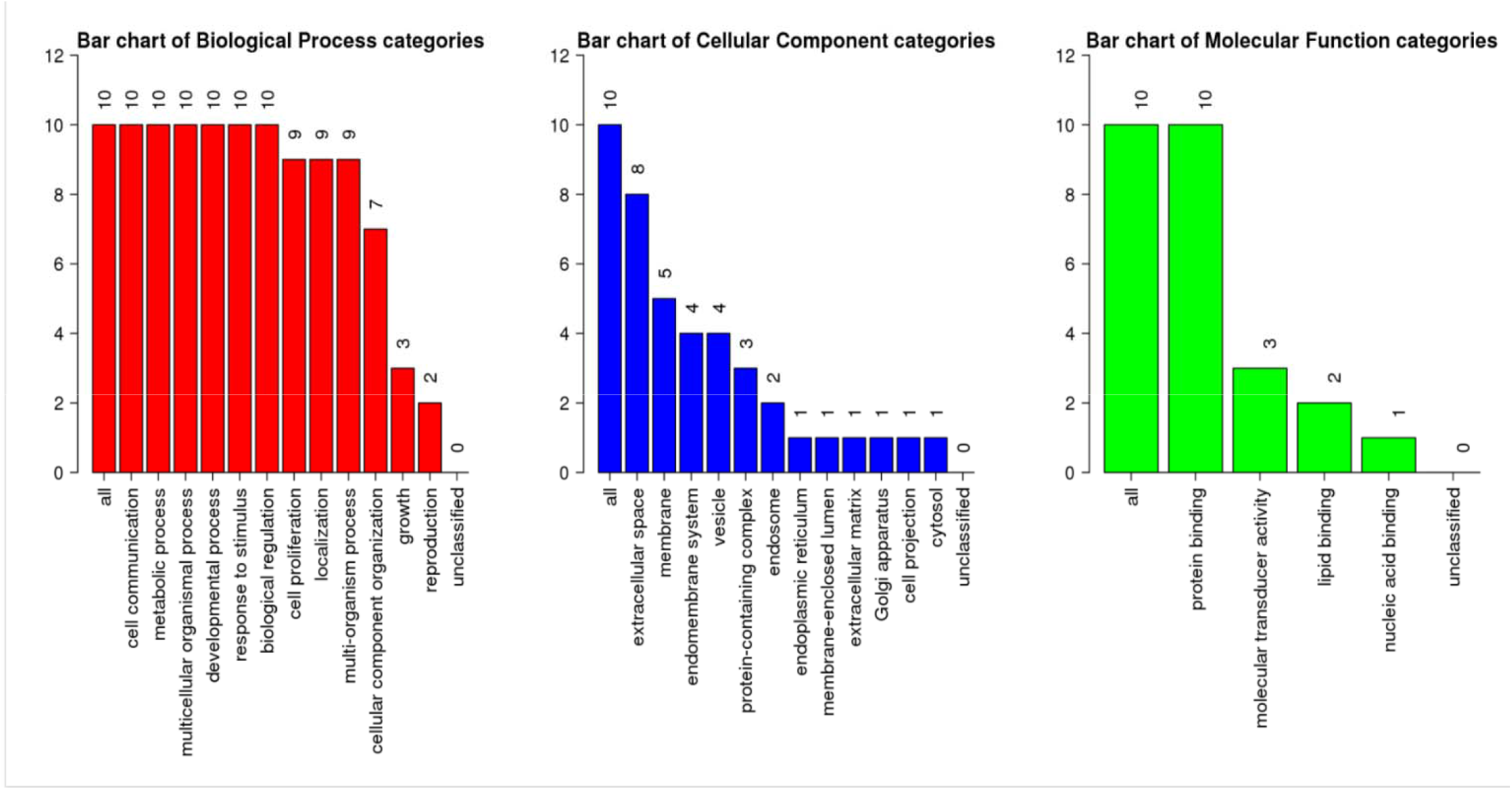
The Gene Ontology of the top 10 common overlapping genes in COVID-19 and COPD.

**Figure 4.**
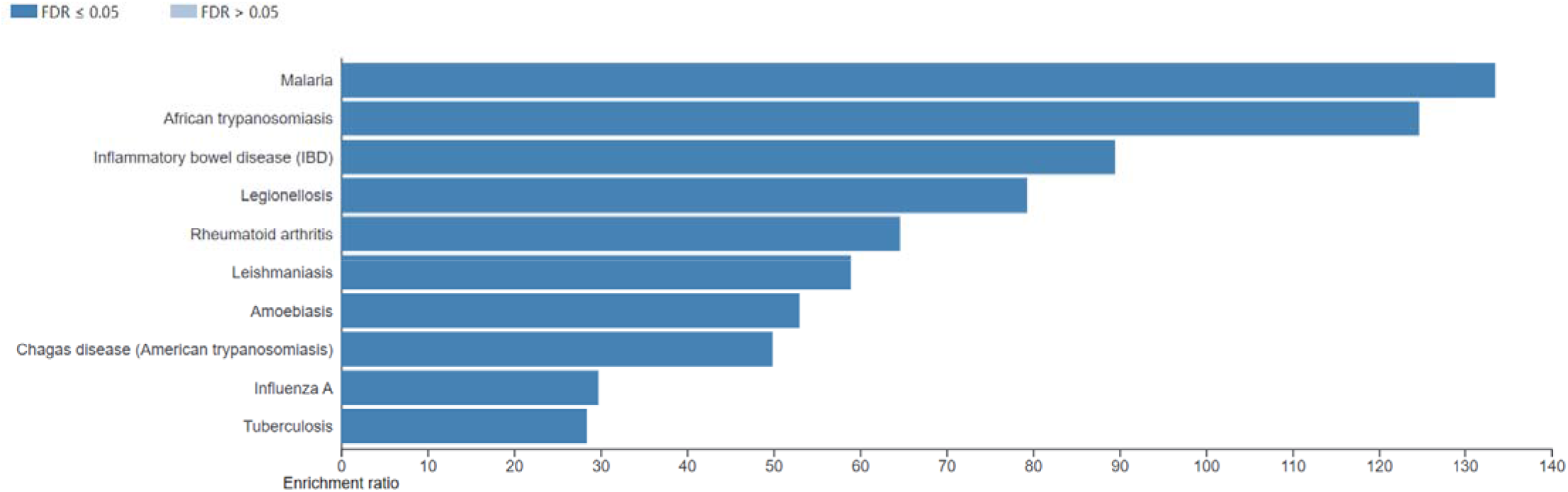
The Gene Pathway analysis of the top 10 common overlapping genes in COVID-19 and COPD.

In **figure 3** for the biological process categories, it can be seen that the highest percentage of the top 10 common overlapping genes are involved in cell communication, metabolic process, multicellular organismal process, development process, response to stimulus and biological regulation, and the lowest percentage of these genes are involved in the reproduction process. While for the cellular components categories, it can be seen that the highest percentage of these genes are found at all parts and also the extracellular space while the least percentage is found to be at the cytosol region. For the molecular functions categories, the highest percentage of these genes are involved in all types of functions and also involved in protein binding while the lowest percentage of these genes are involved in nucleic acid-binding. Besides that, **figure 4** which shows the gene pathway for these top 10 common overlapping genes, shows that most of these genes are involved in the malaria pathway, African trypanosomiasis pathway, inflammatory bowel disease (IBD) pathway, and Legionellosis pathway. While some of these genes are also involved in the Rheumatoid arthritis pathway, Leishmaniasis pathway, Amoebiasis pathway, and Chagas disease (American trypanosomiasis) pathway. The pathway that has been least used by these genes are influenza A pathway and the tuberculosis pathway.

The first common overlapping gene which is also known as the hub gene is IL10. IL10 which is also known as Interleukin 10 is a well-known potential regulatory cytokine that associates the decrease in the inflammatory response. Studies show that the low level of IL10 in the airway tract enhances the development of severe airway inflammation in COPD [27]. The COVID-19 patients who were under ICU support can be easily differentiated with the lower level of the IL10 gene [28]. The level of IL10 also effectively changes for the treatment and progression [29]. In some cases, it is also said that IL10 can be a marking for the disease severity [30]. Even though little research on this molecule has been done, growing evidence supports its significance. Because IL-10, the major member of the IL-10 superfamily, is necessary for the resolution of peripheral inflammation, it has received a great deal of attention as an anti-inflammatory cytokine. In severely/critically ill people, a substantial increase in interleukin IL-10 levels has been found as a crucial feature of COVID-19. Patients with COVID-19 admitted to the intensive care unit (ICU) had considerably greater systemic IL-10 levels than non-ICU patients. Increased IL-10 levels are significantly associated with increased IL-6 and C-reactive protein levels. This suggests that IL-10 may play a role in the severity of COVID-19. IL-10 plays a critical role in chronic airway inflammatory diseases, and individual differences in IL-10 may contribute to disease risk. The serum levels of IL-10 correlated with the clinical factors indicating that they can serve as indicators to estimate the progression of COPD and COVID-19. Previously published preliminary findings suggest that IL-10 may have a role in the severity of SARS-CoV-2-associated sickness and ARDS by raising ACE2 expression and that IL-10 stimulation may have therapeutic potential [31].

The second transcription factor that acts as a hub gene is TLR4. Both innate and adaptive immune cells express TLR4 (neutrophils, macrophages, dendritic cells, endothelial cells, epithelial cells of the skin, and mucous membranes). TLR4 is necessary for the start of inflammatory responses, and overstimulating it can lead to hyperinflammation. SARS-CoV-2’s proteins bind to the ACE2 receptor on host cells, causing membrane fusion and viral RNA release. Following this, viral genomic RNA or dsRNA intermediates are recognised as pathogen-associated molecular patterns (PAMPs) by PRRs, particularly RIG-I-like receptors (RLRs) and Toll-like receptors (TLRs), which drive the relevant downstream signalling cascades required for virus control and eradication. TLRs are important PRRs for recognising viral components or replication intermediates during infection. TLR3 detects viral nucleic acid in the endosome, whereas TLR2 and TLR4 detect viral proteins on the cell surface. TLR4 is also known as toll-like receptor 4 is indeed a major aspect of respiratory epithelial layer host defense. Extreme COPD is linked to decreased expression of TLR4 relative to less serious disorder, with a strong association between expression of the nasal and tracheal [32]. TLR4 may recognize the virus’ structural and non-structural proteins and the resulting development and secretion of cytokines. This is located on the cell wall and is activated by viral glycoproteins [33]. Compensation impacts of both the identification paths needed to excite the antiviral immune system may, ironically, contribute to an inflammatory environment that promotes secondary infections. TLR4 exhibits the highest protein-protein interaction with the SARS-CoV-2 spike glycoprotein, relative to other TLRs. In an immunopathogenic environment, SARS-COV-2 significantly increases interferon-stimulated gene (ISG) expression in the respiratory tract. The fact that ISG activation results in increased ACE2 expression and that pulmonary surfactants in the lung inhibit viral infection by binding to and activating TLR4 suggests that the virus may be binding to and activating TLR4 in order to increase ACE2 expression, which results in increased viral entry. This is supported by the fact that ISGs occur downstream of TLR4-interferon signalling, establishing a link between the two occurrences [34].

The third hub gene is Tumor necrosis factor (TNF). TNF is also known as tumor necrosis factor which involves systemic inflammation. In COPD cases, TNF secretion by pulmonary epithelial cells and sputum cells was greater [35]. Several previous investigations have shown that some pro-inflammatory cytokines play a critical role during acute lung damage, such as acute pancreatitis and sepsis. Additionally, they revealed that infection with viral agents increases the production of cytokines such as Tumor Necrosis Factor alpha (TNF-alpha), which is a key mediator of inflammation. TNF appears in the bloodstream and infection tissues of COVID-19 patients, and TNF is essential in virtually all chronic inflammation reactions, serving as an inflammatory amplification [36]. It is found in a study that COPD patients with acute COVID□19, had increased TNF levels [37].

The fourth hub gene is IL6 which is known as Interleukin 6 is a pro-inflammatory cytokine and immunomodulatory pleiotropic cytokine naturally produced by epithelial airway cells, alveolar macrophages, adipocytes, and myocytes, and much other body tissue. They make strong of IL6 in the pathogenicity of chronic obstructive pulmonary disease (COPD) is recommended by research proving that high serum or sputum IL6 levels are linked to impaired lung function or a faster decrease in lung function. IL6 has been correlated with skeletal muscle weakness in COPD, as well as aggravate and pulmonary infections in COPD patients [38]. IL6 is an effective agent of inflammatory and has proved to be an important indicator of extreme COVID□19 in studies. IL6 is liable for the altitude and suppression of albumin synthesis of acute-phase reactants such as hepcidin, fibrinogen, serum amyloid A, and C□reactive protein. Autoimmunity and chronic inflammation have been directly linked to the disordered production of IL6 [39]. Consequent swelling of the lungs in COVID-19 indicates aggravation in COPD patients is probable [40].

The fifth hub gene is CXCL8, which is also known as Interleukin 8. During a severe aggravation of COPD, the respiratory muscles weakness is increased. CXCL8 acts as a trafficking mediator for neutrophils. This chemokine is highly involved in inflammatory processes especially those associated with viral infections. CXCL8 has been engaged in creating respiratory muscle weakness in both hospitalized and sustainable COPD patients [41]. CXCL8 is further increased during exacerbations of COPD. The production of a huge amount of chemokine CXCL8 is also reported as the main cause of death in COVID-19 patients [14]. A systematic statistical study mentioned that CXCL8 includes the lung problems of COVID-19 high-risk patients as the main genes [42].

The sixth hub gene present is IL4 known as Interleukin 4 plays an eminent role in the activation of the JAK-STAT pathway and leads to the disturbing pro-inflammatory cytokine. It is engaged in hyper responsibility (AHR) for the airways. AHR is among COPD symptoms [43]. Specific elements of type 2 inflammatory process, such as type 2 cytokine IL4 and eosinophil aggregation, could potentially have beneficial effects against COVID-19 [44].

The seventh hub gene present is ICAM1 is known as intercellular adhesion molecule 1. The first stage in the inflammation cycle would be over-production of the adhesion molecules, resulting in premature neutrophil liberation. Another of these adhesion molecules is ICAM1 which is increased in patients with COPD [45]. Levels of ICAM1 increased in patients with a moderate illness, increased significantly in serious conditions, and reduced in the process of convalescence. This can be inferred that the elevated expression of intracellular cell adhesion molecules is related to the extent of COVID-19 disease and can lead to defective coagulation [46]. Most preprint reports now exist that reveal SARS□CoV-2 can attach four various intracellular binding sites in humans. Most of these seem so to be highly expressed in COPD, as notable in commonly diagnosed pathogen adhesive proteins such as ICAM1 have been found [47].

IFNG is the eight hub gene present and it is also known as Interferon-gamma. COPD patients are conditioned to a high, more than five times greater chance of serious COVID-19 infection. Potentially angiotensin approach that enables 2 (ACE2) is the binding factor between COPD former existence and COVID-19 frequency. An operating system and platform review identified IFNG as the top core genes for high-risk group COVID-19 patients in lung disease [42].

TLR2, or toll-like receptor 2, is the ninth hub gene involved. TLR2 expression is connected with lung function level and decline, as well as the frequency of inflammatory processes in the sputum in patients with COPD, suggesting a role in the development and progression of COPD [48, 49]. While coronavirus virulence pathogenesis is developing, TLR2 is now known to identify virus particles and could therefore possibly be engaged in the cell pathways contributing to pulmonary embolism of COVID-19 [50].

The final hub gene involved is IL18 alias, Interleukin 18. The research provides good support to IL18 being implicated in COPD pathophysiology. For preclinical models of inflammation and damage, the neutralization of antibodies to IL18 has an effect. In COPD, IL18 is viewed as a possible potential treatment-limiting disintegration and redoing [51]. Whereas COVID-19 virulence factors are complicated, a knowledge of the process of inflammatory response in this disease is very important. As a direct consequence of viruses’ stimulation of cytoplasmic inflammasome NLRP3, the virus and pathogenesis correlated with SARS coronaviruses evolve. This inflammasome produces pro-inflammatory cytokine IL18 within macrophages and Th1 lymphocytes that also dictates the virulence inflammatory response accountable for the infectiousness and illnesses of COVID-19 [52]. In summary, COPD patients with severe COVID-19 showed higher serum levels of inflammatory cytokines such as IL-2, IL-6, IL-8, IL-10, and TNF-α, indicating that the immune response may be altered globally.

## 4) CONCLUSION

Our study found that both COVID-19 and COPD share hub genes based on protein-protein interaction analysis. These hub genes and enrichment may have significant clinical implications for COPD patients who are at an increased risk of developing severe illness because of COVID-19.

## Notes

### Competing Interest Statement

The authors have declared no competing interest.

